# Clustering-based time series analysis on insulin response in the blood-brain barrier

**DOI:** 10.1101/2021.03.07.434315

**Authors:** Krishna R. Kalari, Zengtao Wang, Xiaojia Tang, Suresh K. Swaminathan, Karunya K. Kandimalla

**Affiliations:** Department of Health Sciences Research, Mayo Clinic, 200 First Street SW, Rochester, MN USA; Department of Pharmaceutics and Brain Barriers Research Center, University of Minnesota, MN, USA

**Keywords:** Insulin signaling, Time-series analysis, Longitudinal data, Blood-brain barrier, Transcriptomics, Molecular pathways

## Abstract

**Background:** Critical functions of the blood-brain barrier (BBB), including cerebral blood flow and vascular response, are regulated by insulin signaling pathways. Therefore, endothelial insulin resistance could lead to vascular dysfunction, which is associated with neurodegenerative diseases such as Alzheimer’s disease (AD).

**Objective:** The objective of the current study is to map the dynamics of insulin-responsive pathways in polarized human cerebral microvascular endothelial cells (hCMEC/D3) cell monolayers, a widely used BBB cell culture model, to identify molecular mechanisms underlying BBB dysfunction in AD.

**Methods:** RNA-Sequencing (RNA-Seq) was performed on hCMEC/D3 cell monolayers with and without insulin treatment at various time points. The Short Time-series Expression Miner (STEM) method was used to identify clusters of genes with distinct and representative patterns. Functional annotation and pathway analysis of the genes from top clusters were conducted using the Webgestalt and Ingenuity Pathway Analysis (IPA) software, respectively.

**Results:** Quantitative expression differences of 19,971 genes between the insulin-treated and control monolayers at five-time points were determined. STEM software identified 11 clusters with 3061 genes across that displayed various temporal patterns. Gene ontology enrichment analysis performed using the top 5 clusters demonstrated that these genes were enriched in various biological processes associated with AD pathophysiology. The IPA analyses revealed that signaling pathways exacerbating AD pathology such as inflammation were downregulated after insulin treatment (clusters 1 to 3). In contrast, pathways attenuating AD pathology were upregulated, including synaptogenesis and BBB repairment (clusters 4 and 5).

**Conclusions:** These findings unravel the dynamics of insulin action on the BBB endothelium and inform about downstream signaling cascades that potentially regulate neurovascular unit (NVU) functions that are disrupted in AD.

## INTRODUCTION

Cerebrovascular endothelium, commonly referred to as the blood-brain barrier (BBB), is instrumental in maintaining vascular response to regulate cerebral blood flow, delivering essential nutrients for sustaining brain functions and removing toxic metabolites from the brain [1]. In addition, the BBB serves as a formidable barrier protecting the brain from circulating xenobiotics and immune challenges emanating from the periphery [2]. The BBB accomplishes these diverse functions by not functioning independently but as a crucial part of the neurovascular unit (NVU), which is organized by the precise spatial arrangement and well-coordinated communication among various cells in the cerebral vasculature (endothelial cells, pericytes, and smooth muscle cells) and brain parenchyma (astrocytes and neurons) [3]. The molecular mechanisms regulating NVU composition and function in health and disease are only partially understood because of the paucity of molecular-level information on the less abundant yet functionally critical cerebrovascular endothelial cells and pericytes.

Studies have shown that BBB dysfunction is associated with neurodegenerative diseases such as Alzheimer’s disease (AD) and Parkinson’s disease [4, 5]. Recent research has revealed the importance of hyperinsulinemia and peripheral insulin resistance in AD pathogenesis. As several important BBB functions were shown to be handled by insulin signaling pathways [6], endothelial insulin resistance could compromise BBB integrity and function, and lead to BBB dysfunction. Therefore, it is crucial to study insulin’s impact and characterize insulin responsive pathways in the BBB endothelium. RNA-Sequencing technology represents a powerful tool to investigate transcriptomic changes across the genome and detect insulin’s effect on gene regulation using model systems.

Given the impact of insulin on blood-brain-barrier endothelial cells, we have performed RNA-Sequencing (RNA-Seq) of polarized human cerebral microvascular endothelial cells (hCMEC/D3) cell monolayers with and without insulin treatment at various time points to identify molecular and cellular pathways that are responsive to insulin. We conducted experiments at five time points to differentiate insulin-responsive pathways and identify early and late response molecular and cellular pathways activated or inhibited after insulin treatment in the hCMEC/D3 cell monolayers. To our knowledge, this is the first systematic study investigating the transcriptional response to insulin in BBB endothelial cells in a time series RNA-Seq experiment.

## METHODS

### Cell Culture and Illumina TruSeq v2 mRNA Protocol

Immortalized human cerebral microvascular endothelial cell line (hCMEC/D3) was kindly provided by P-O Couraud, Institut Cochin, France. The polarized endothelial monolayers were cultured as described previously [7], and the detailed methods are provided in our previous publication [8]. In this study, we focused on the hCMEC/D3 cell lines that were treated with 100 nm insulin at various time points (t=10, 20, 40, 80, and 300 minutes). We also harvested synchronized control BBB cell monolayers without insulin treatment at each time point (as shown in **Figure 1**). Paired-end RNA libraries from insulin-treated and control hCMEC/D3 cell monolayers (a total of 10 samples with 100mg insulin at five time points) were prepared according to the manufacturer’s instructions using TruSeq RNA Sample Prep Kit v2 (Illumina, San Diego, CA). A detailed protocol of the RNA library preparation has been discussed in our previous publication [8].

**Figure 1:**
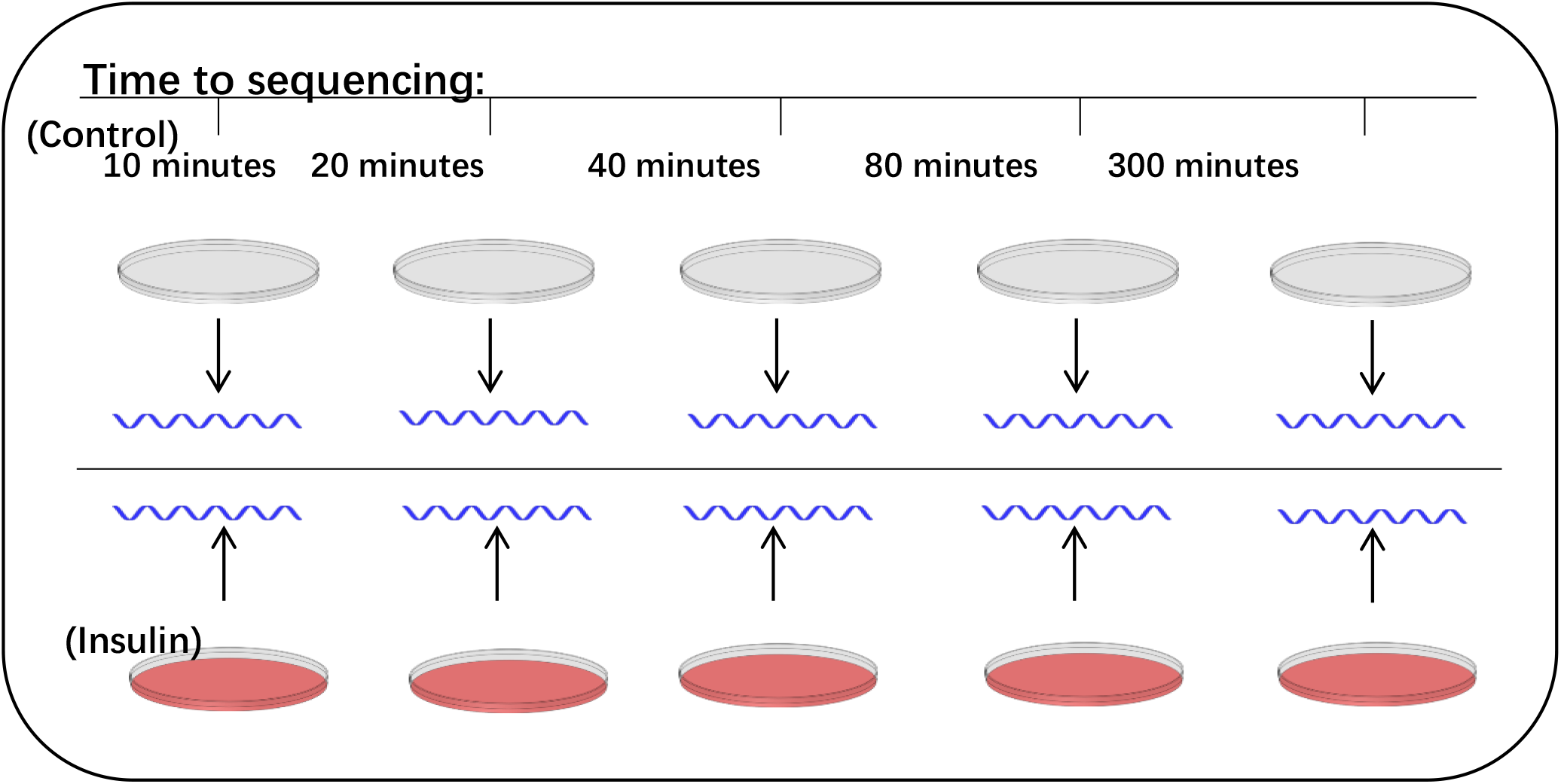
Experimental design showing the RNA-Sequencing time-points of human cerebral microvascular endothelial cell (hCMEC/D3) monolayers with and without insulin treatment.

### RNA-Seq Data Processing and gene expression quantification

After paired-end transcriptome sequencing, the RNA-Seq data of ten samples were processed using the MAP-RSeq — a comprehensive computational workflow developed at the Mayo Clinic to obtain various genomic features from the RNA-Seq experiment [9]. The main goal of the MAP-RSeq pipeline is to obtain multiple genomic features, such as gene expression, exon counts, fusion transcripts from RNA-Seq data. MAP-RSeq provides quality control reports and summary statistics of the sequencing reads. Reads were mapped to the human genome reference hg19 build, and the total number of reads, mapped reads, the number of reads mapped to the genome, and the numbers of reads mapped to junctions were obtained for each sample. Gene expression counts were quantified using the HT-Seq module (http://www-huber.embl.de/users/anders/HTSeq/doc/count.html) from MAP-RSeq pipeline for five hCMEC/D3 controls and five matched insulin-treated hCMEC/D3 monolayers. Principal component analysis and data analysis were then conducted using R programming language and the gene expression data obtained from these five treatment and control pairs.

### Time-series Gene Expression Analysis

Normalized gene expression data was used for time-series data analysis. We determined the difference between the paired samples (insulin-treated versus control monolayers) at all the five time points for every gene. The resulting gene expression differences were then provided as input to the Short Time-series Expression Miner (STEM) [10] method to identify clusters of genes with distinct and representative patterns. STEM is an application specifically designed for the clustering analysis of short time-series gene expression data (3-8 time points). The STEM method identifies clusters of genes that are statistically significant based on the correlation coefficient (the minimal default correlation is 0.7). In the STEM software, we chose the parameter options such as the maximum number of model profile=50, a maximum unit change in model profiles between time points = 2, and clustering method= STEM. The significance of a cluster is determined by STEM software based on a binomial distribution by comparing the actual number of genes assigned to the group against the expected number of genes.

### Gene set and pathway analysis

The top clusters of genes identified by the STEM software were retrieved and used to conduct overrepresentation analysis using the webGestalt software and gene ontology database [11]. Pathway analysis of genes from the significant clusters was performed using Ingenuity pathway analysis software (QIAGEN Inc., https://www.qiagenbioinformatics.com/products/ingenuitypathway-analysis) to identify pathways that are activated or predicted to be inhibited.

## RESULTS

Due to the cost of RNA-Sequencing and lack of availability of tissue samples, most of the gene expression studies are single snap-shot studies to identify differentially expressed genes/pathways between two conditions. Biological processes are often dynamic and require temporal monitoring to decipher their response to health and disease. Insulin signaling pathways that are characterized by rapid response upon stimulation and quick return to the baseline levels are prime examples of the pathways that require dynamic monitoring to capture pathophysiological variations by conducting time-series experiments. However, it is challenging to obtain serial samples from the tissue of an individual, particularly from the brain. Therefore, we have conducted studies using the hCMEC/D3 monolayers to study insulin treatment dynamics in the blood-brain barrier. At every time point, gene expression changes in insulin-responsive pathways are investigated from the paired data of control and insulin-treated hCMEC/D3 monolayers. RNA libraries were prepared and sequenced as described in the methods section. After removing the control and treatment samples that were not sequenced in the same batch, a total of ten samples remained for further time-series data analysis. Raw gene expression counts were obtained, and the total read depth across samples varied from 92-149 million reads. Of the 23,399 genes with expression data, there were 3428 genes with less than 32 counts across all ten samples. After filtering out the genes with zero counts, we had 19,971 genes for the rest of the time-series data analysis. We determined the quantitative gene expression differences between the treatment and control at all five time points (t=10, 20, 40, 80, and 300 minutes). We analyzed the data with short-time-series data analysis software (STEM).

### Time-series data analysis identifies eleven significant gene clusters in hCMEC/D3 monolayers after insulin treatment

After removing the low expressed genes in this study, we have provided the normalized time-series gene expression data as the input to the STEM software [10]. The STEM method assigned all the genes to a set of pre-defined temporal expression profile patterns defined by the number of time points and maximum units of fold change. The STEM clustering method represents a set of distinct and representative model temporal expression profiles independent of the input data; these model profiles correspond to possible profiles in the input gene expression data. All of the model profiles begin at zero, and the model profile is expected to increase, decrease, or remain stable (do not change) between two-time points. As shown in **Figure 2**, STEM software identified 11 clusters with 3061 genes across the five time points that display various patterns. The number of genes in each cluster is indicated in **Table 1**. For each cluster, the p-value is determined based on a permutation test by comparing the number of genes assigned to a particular profile and genes that are expected to follow that specific profile by chance (as shown in **Figure 2)**. After a thorough investigation of the individual clusters, we have observed that clusters 1, 2 and 3 (2096 genes), as well as clusters 4 and 5 (234 genes), followed similar temporal expression profiles (**Figure 2**). The gene expression in clusters 1, 2 and 3 went down at 40 minutes after insulin treatment, and the gene expression in clusters 4 and 5 went up after insulin treatment. Hence, we compared the genes and pathways represented by these clusters that demonstrated contrasting trends.

**Figure 2:**
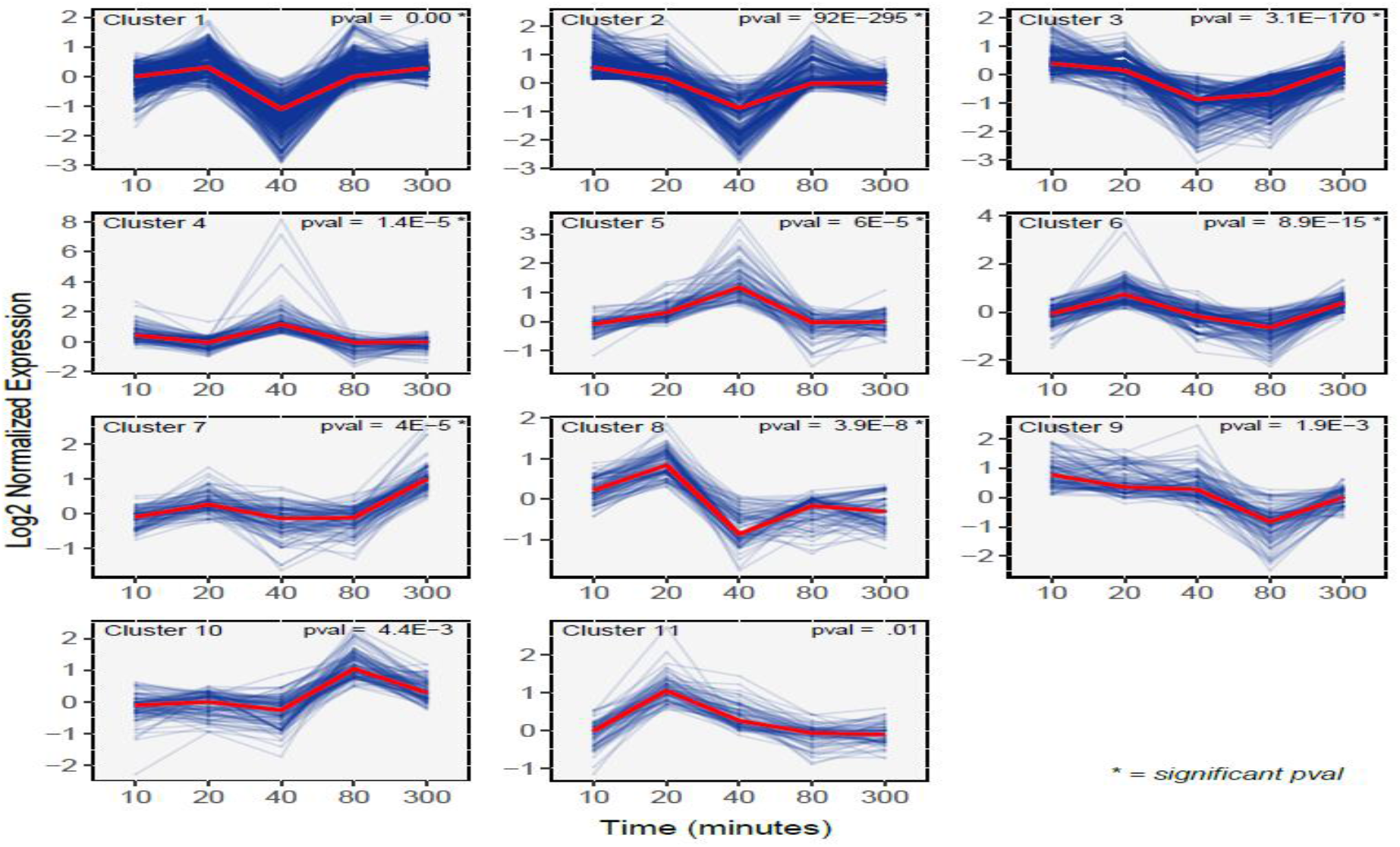
The figure shows the eleven clusters or profiles of genes after insulin treatment (treatment –control gene expression data) from five-time points 10 minutes, 20 minutes, 40 minutes, 80 minutes, and 300 minutes. The clusters are ordered based on the number of genes ranked by significance. Inside each set (in the box), on the top left-hand corner, the cluster-ID is indicated, and on the right-hand corner, the enrichment p-value is displayed.

**Table 1:** Time series clustering of the genes based on gene expression profile changes was conducted using the STEM software. STEM detected 11 significant clusters of genes with similar expression profiles across five-time points. The number of genes in each cluster is listed.

| Time Series |  |
| --- | --- |
| Data | Number of Genes |
| Cluster 1 | 926 |
| Cluster 2 | 701 |
| Cluster 3 | 469 |
| Cluster 4 | 135 |
| Cluster 5 | 99 |
| Cluster 6 | 174 |
| Cluster 7 | 119 |
| Cluster 8 | 112 |
| Cluster 9 | 124 |
| Cluster 10 | 120 |
| Cluster 11 | 82 |

### Functional annotation of genes in the top 5 clusters affected by insulin treatment shows the genesets involved in AD pathophysiology

We obtained the genes from the top five clusters (cluster 1-3=2096 and cluster 4-5 = 234 genes) and conducted functional annotation of the genes using the Webgestalt software. Gene ontology (GO) enrichment analysis was performed using the 2096 geneset (clusters 1-3) that were downregulated at a 40-minute time-point after insulin treatment. The genes are highly enriched in various biological processes such as the GPCR signaling genes, neuropeptide signaling genes, coupled to cyclic nucleotide second messenger genes, peptide cross-linking genes, and humoral immune response genes **(Table 2)**. Similarly, the GO enrichment analysis was conducted for 234 geneset (clusters 4&5) that were upregulated at 40 min time frame following insulin treatment in the hCMEC/D3 monolayers (**Table 3**). These genes are associated with biological processes that are also associated with AD pathophysiology, such as central nervous system neuron differentiation, response to mechanical stimulus, skeletal system morphogenesis, extracellular structure organization, and mesenchymal cell proliferation genes, connective tissue development, embryonic skeletal system development.

**Table 2:**
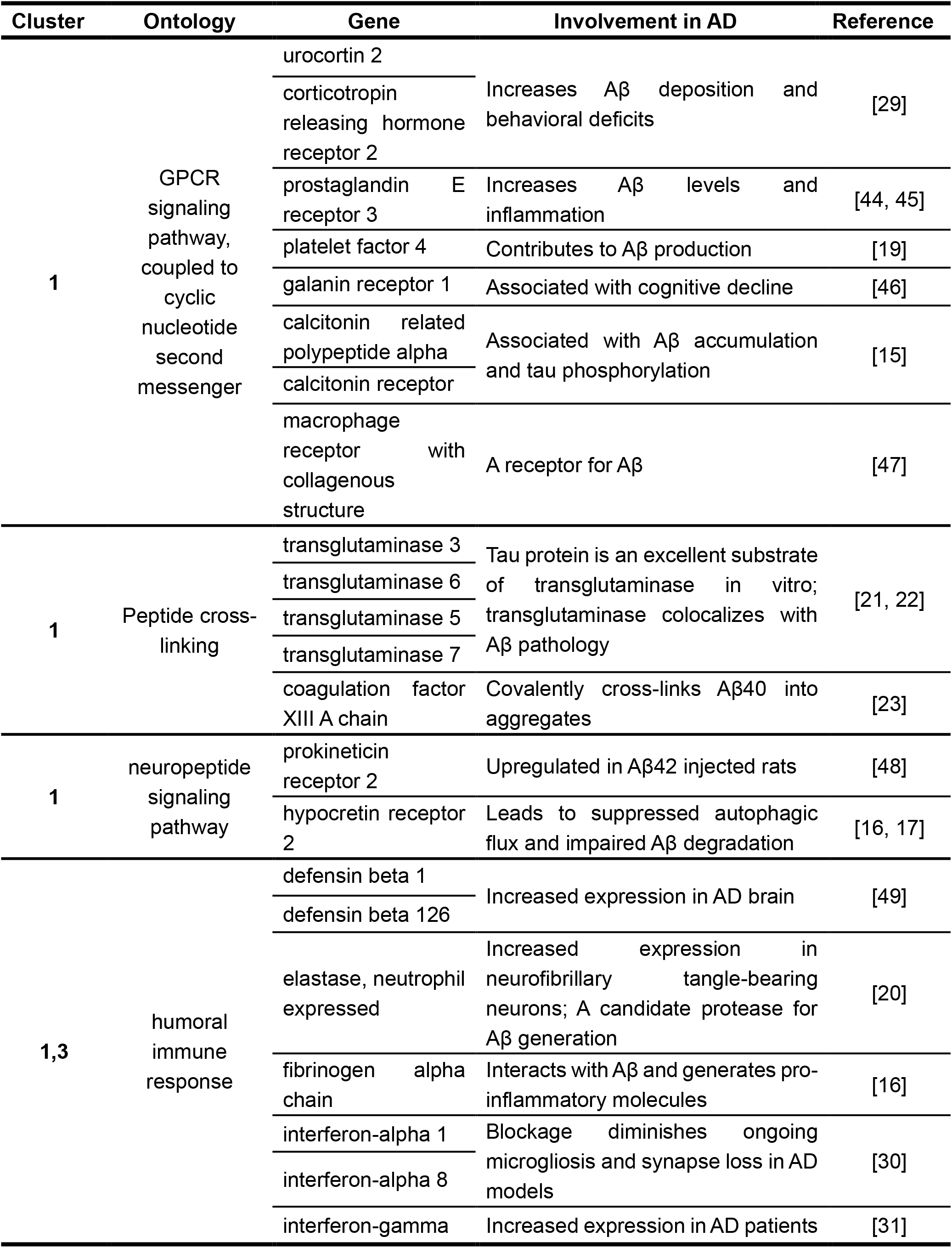
Involvement of cluster 1-3 genes in AD pathology after GO enrichment analysis. These genes are downregulated at 40 minutes after insulin treatment.

**Table 3:** Involvement of cluster 4-5 genes in AD pathology after GO enrichment analysis. These genes are upregulated at 40 minutes after insulin treatment.

| Cluster | Ontology | Gene | Involvement in AD | Reference |
| --- | --- | --- | --- | --- |
| 4 | mesenchymal cell proliferation | bone morphogenetic protein 7 | Prevented neuronal injuries in PC12 cells induced by Aβ25-35 | [32] |
| 4 | embryonic skeletal system development | platelet derived growth factor receptor alpha | defective PDGF-BB-PDGFR-β signaling resulted in faulty amyloid-β clearance | [18] |
|  |  | bone morphogenetic protein 7 | Discussed above |  |
| 4 | connective tissue development | collagen type VI alpha 3 chain | Treatment of neurons with soluble collagen VI blocked the association of Aβ oligomers with neurons and neurotoxicity | [24] |
|  |  | collagen type VI alpha 1 chain |  |  |
|  |  | collagen type VI alpha 2 chain |  |  |
| 4 | central nervous system neuron differentiation | plexin A4 | Decreased expression in AD brain | [50] |
|  |  | dopamine receptor D2 | Decreased expression in AD brain; suppresses neuroinflammation | [33, 34] |
| 4 | response to mechanical stimulus | Decorin | Decreased expression in AD brain | [51] |
|  |  | dopamine receptor D2 | Discussed above |  |
| 5 | skeletal system morphogenesis | bone morphogenetic protein 7 | Discussed above |  |
|  |  | collagen type VI alpha 2 chain | Discussed above |  |
|  |  | collagen type VI alpha 3 chain | Discussed above |  |
|  |  | platelet derived growth factor receptor alpha | Discussed above |  |
|  |  | collagen type VI alpha 1 chain | Discussed above |  |
| 5 | extracellular structure organization | Decorin | Discussed above |  |
|  |  | bone morphogenetic protein 7 | Discussed above |  |
|  |  | collagen type VI alpha 2 chain | Discussed above |  |
|  |  | collagen type VI alpha 3 chain | Discussed above |  |
|  |  | collagen type VI alpha 1 chain | Discussed above |  |

### Canonical pathway analysis of the top five clusters has identified pathways that are critical for NVU functions

We conducted pathway analysis to identify the known canonical pathways that are associated with our top clusters. The 2096 geneset (genes from cluster 1-3) and 234 geneset (genes from clusters 4-5) were submitted to the Ingenuity Pathway Analysis (IPA software), and the pathways with a p-value < 0.05 were obtained. The top canonical pathways are shown in Table 4 and Table 5 with the z-scores; if the z-score > 2, the pathway is deemed to be activated after insulin treatment, and if Z-score < −2, the pathway is predicted to be inhibited after insulin treatment. The ratio column in Tables 4 and 5 indicates the proportion of genes from our geneset to the genes that are present in the canonical pathway.

**Table 4.** Ingenuity pathway analysis of the genes from clusters 1, 2, and 3. List of pathways, with z-score, −log(p-values), and the ratio are listed below.

| **Ingenuity Canonical Pathways** | **z-score** | **-log(p-value)** | **Ratio** |
| --- | --- | --- | --- |
| SPINK1 Pancreatic Cancer Pathway | 2.530 | 5.740 | 0.183 |
| CREB Signaling in Neurons | -6.633 | 7.510 | 0.076 |
| Breast Cancer Regulation by Stathmin1 | -5.692 | 5.560 | 0.068 |
| Neuroprotective Role of THOP1 in Alzheimer's Disease | -3.162 | 3.630 | 0.103 |
| Intrinsic Prothrombin Activation Pathway | -2.449 | 3.580 | 0.167 |
| Amyotrophic Lateral Sclerosis Signaling | -3.000 | 3.100 | 0.103 |
| MSP-RON Signaling In Macrophages Pathway | -2.828 | 2.600 | 0.089 |
| Neuroinflammation Signaling Pathway | -3.051 | 2.260 | 0.060 |
| MSP-RON Signaling In Cancer Cells Pathway | -3.000 | 2.080 | 0.075 |
| IL-17 Signaling | -3.464 | 1.890 | 0.064 |
| Retinoate Biosynthesis I | -2.000 | 1.710 | 0.118 |
| Role of Pattern Recognition Receptors in Recognition of Bacteria and Viruses | -2.236 | 1.690 | 0.065 |
| eNOS Signaling | -2.449 | 1.600 | 0.063 |
| Nitric Oxide Signaling in the Cardiovascular System | -2.449 | 1.490 | 0.071 |
| Cardiac Hypertrophy Signaling (Enhanced) | -3.900 | 1.470 | 0.046 |
| FGF Signaling | -2.449 | 1.350 | 0.071 |
| Tumor Microenvironment Pathway | -3.162 | 1.340 | 0.057 |
| Netrin Signaling | -2.236 | 1.310 | 0.077 |

**Table 5.** Ingenuity pathway analysis of the genes from clusters 4 and 5. List of pathways, with z-score, −log(p-values) and the ratio are listed below.

| **Ingenuity Canonical Pathways** | **z-score** | **-log(p-value)** | **Ratio** |
| --- | --- | --- | --- |
| GP6 Signaling Pathway | 2.236 | 2.940 | 0.040 |
| White Adipose Tissue Browning Pathway | 2.000 | 2.880 | 0.039 |
| IL-6 Signaling | 2.000 | 2.090 | 0.032 |
| Dendritic Cell Maturation | 2.000 | 1.540 | 0.022 |
| Synaptogenesis Signaling Pathway | 2.236 | 1.330 | 0.016 |

## DISCUSSION

The BBB is a critical gatekeeper that dynamically removes toxic metabolites from the brain and delivers essential nutrients, such as glucose and insulin, to maintain brain functions. BBB dysfunctions manifested as reduced cerebral blood flow, impaired amyloid-beta clearance, and cerebrovascular inflammation are believed to promote AD progression. Notably, these pathological features are consistently indicated in type-2 diabetes mellitus (T2DM) and are associated with insulin resistance, which is an established risk factor for AD [12, 13]. However, the cause-and-effect relationship between cerebrovascular dysfunctions and insulin resistance in AD is poorly characterized. Therefore, it is imperative to identify insulin-responsive pathways in the BBB endothelium, a major component of BBB.

### GO enrichment analysis

One of the major AD pathologies is abnormal amyloid-beta (Aβ) deposition in the brain, which is due to the imbalance between production and its clearance from the brain. Peripheral hyperinsulinemia has been shown to increase brain and plasma Aβ levels significantly [33]. Our previous study further demonstrated insulin could differentially impact BBB trafficking of Aβ40 versus Aβ42 [14]. However, the underlying molecular mechanisms are not well understood. In the current study, gene expression changes in polarized hCMEC/D3 monolayers upon insulin treatment were clustered by STEM analysis. In clusters 1-3, the gene expression decreased at 40 minutes post insulin treatment, whereas the opposite trend was observed in clusters 4 and 5. Notably, genes identified in clusters 1-3 were shown to upregulate Aβ levels, whereas genes from clusters 4 and 5 were reported to decrease Aβ levels. For instance, calcitonin-related polypeptide and calcitonin receptor were found in clusters 1-3, and treatment with calcitonin gene-related peptide (cGRP) receptor antagonists could reduce Aβ accumulation [15]. Hypocretin receptor 2 (identified in clusters 1-3), which binds to neuropeptides orexin A and orexin B, was found to be downregulated after insulin treatment. Orexin was demonstrated to disrupt the autophagosome-lysosome fusion process and may lead to impairment in Aβ degradation [16]. Further, the amount of brain Aβ was found to be substantially decreased upon knocking down the orexin gene [17]. In contrast, platelet-derived growth factor receptor, which mediates Aβ clearance by LRP1 [18], was identified in clusters 4-5, and the expression was increased with insulin treatment. On the other hand, platelet factor 4 (PF4) and elastase, which could potentially enhance Aβ production, were identified in clusters 1-3. While PF4 is released from activated platelets which produce more than 90% of circulating Aβ [19], elastase is a proteolytic enzyme that is a candidate protease involved in the generation of Aβ [20]. Apart from production and clearance, genes that regulate Aβ aggregation were also segregated in clusters 1-3. For example, several isoforms of transglutaminase, associated with peptide cross-linking, were found in clusters 1-3. Transglutaminase is a calcium-activated enzyme that converts soluble proteins into insoluble species through cross-linking. Transglutaminase levels and activity are increased in AD brains [21]. Further, Aβ has been shown to be a substrate of transglutaminase [22]. Therefore, it is possible insulin could mitigate Aβ deposition by modulating the transglutaminase expression. Coagulation factor XIII, another enzyme that belongs to the same geneset, has been shown to covalently cross-link Aβ40 into oligomers as well as fibrins [23].

On the other hand, genes that reduce Aβ aggregation were identified in clusters 4-5. For example, three isoforms of collagen type Ⅵ alpha chain were found to increase after insulin treatment. Reduction of collagen Ⅵ could enhance Aβ aggregation and prevent neurotoxicity. Further, treatment of soluble collagen Ⅵ was shown to block the association of Aβ oligomers with neurons [24].

Inflammation, which is mediated by diverse immune cells and pro-inflammatory chemical signals, is a common pathological feature of both AD and diabetes [25]. Insulin has been shown to prevent inflammation in both peripheral tissues and the brain [26–28]. However, the anti-inflammatory effect of insulin on BBB endothelial cells is not well understood. Genes from clusters 1-3 that are downregulated after insulin treatment were found to drive inflammation, whereas genes in clusters 4 and 5 could reduce inflammation. For example, humoral immune response gene ontology was significantly enriched in clusters 1-3, and many genes were involved in pro-inflammatory actions, one of which is the fibrinogen alpha chain that encodes the alpha subunit of fibrinogen. Aβ was shown to interact with fibrinogen and lead to the production of pro-inflammatory molecules [16]. Moreover, both urocortin 2, which is a member of corticotropin-releasing factor (CRF), and corticotropin-releasing hormone receptor 2 were found in clusters 1-3. Overexpression of brain CRF was shown to mimic the chronic inflammatory stress and could further increase Aβ deposition and behavioral deficits [29]. Several isoforms of interferon (INF) were also in this geneset, including type Ⅰ (INF-α 1, INF-α 8) and type Ⅱ class (INF-γ). Previous studies have demonstrated that type Ⅰ INF response drives neuroinflammation and synapse loss in AD [30] and higher levels of INF-γ were observed in patients with mild cognitive impairment (MCI) compared to controls [31]. Contrarily, genes in cluster 4-5 were found to have anti-inflammatory effects and upregulated with insulin treatment. Bone morphogenic protein 7 (BMP7) encodes a secreted ligand of the transforming growth factor-beta (TGF-beta) superfamily of proteins. Previous study indicated that BMP7 was able to prevent neuronal injuries induced by Aβ, including cell apoptosis and oxidation stress [32]. Moreover, dopamine receptor D2 was found to be upregulated and astrocytic dopamine D2 receptors were shown to suppress neuroinflammation[33]. Hippocampal dopamine D2 receptor also correlates with memory functions in AD [34].

### IPA canonical pathway analysis

Using Ingenuity Pathway Analysis (IPA) software, we identified several signaling pathways to be enriched in clusters 1-3 and downregulated after insulin treatment. In agreement with GO enrichment analysis, the neuroinflammation signaling pathway was found to be downregulated with insulin treatment. Moreover, specific genes have been reported to be related to AD. For example, matrix metalloproteinase-3 (MMP-3) was shown to distribute in AD brains selectively [35]. Additionally, cognitively healthy individuals with AD risk markers were found to have higher levels of CSF MMP-3 [36]. IL17 pathway was also involved in inflammation and shown to be downregulated after insulin treatment. Previous studies demonstrated that pretreatment with IL-17 neutralizing antibody could markedly reduce Aβ42-induced neurodegeneration and prevent the increase of pro-inflammatory mediators in a dose-dependent manner [37]. Interestingly, treatment with anti-IL-17 antibody was also shown to ameliorate insulin resistance and inflammation in TD2M. Therefore, this pathway could provide yet another mechanistic connection between AD and T2DM. Notably, there are also other pathways that mitigate AD progression were found to be downregulated upon insulin exposure. For instance, endothelial nitric oxide synthase (eNOS) signaling was found to be downregulated. This effect could be induced by excessive insulin exposure that could drive insulin resistance [38].

In clusters 4-5, however, pathways that attenuate AD pathologies were found to be enriched by IPA analysis, one of which is the synaptogenesis signaling pathway. AD is characterized by progressive cognitive impairment and memory loss, which result from loss of hippocampal and cortical synapses. Therefore, one promising AD treatment strategy might be stimulating synaptogenesis. Given that intranasal insulin administration was shown to improve cognition in AD patients, our finding provided a potential mechanism by which insulin exerts such beneficial effects [39]. Although synapse is exclusively formed in neurons, BBB endothelium is essential for synaptogenesis by maintaining the homeostasis of the brain microenvironment and interacting with glia cells and neurons. Li et al. previously showed that activation of endothelial-derived GABA signaling could promote neuronal migration, which is critical for synapse formation [40]. Additionally, several critical genes in this pathway were also identified to be associated with the mitigation of AD pathology, one of which is thrombospondin2 (THBS2). Previous evidence has suggested that thrombospondin1 expression was decreased in AD brains, and treatment of exogenous thrombospondin1 could restore the Aβ induced synaptic pathology [41]. It is possible that thrombospondin2 has similar beneficial effects because these proteins share the same structure and the amino acid sequences are slightly different [42]. GP6 signaling pathway was also enriched in clusters 4-5 and upregulated after insulin treatment. This pathway contains many collagens and laminins that form the basement membrane of the BBB. Particularly, collagen Ⅵ in this pathway was also demonstrated to prevent the neurotoxicity of Aβ [24]. Insulin was previously shown to increase tight junction proteins of BBB endothelial cells [43]; our findings suggested that it could also potentially promote the establishment of the basement membrane. Therefore, it is possible that insulin could restore the BBB disruption, which is highly implicated in AD progression. In summary, these results provided novel insights into the molecular mechanisms by which insulin exerts beneficial effects on the BBB endothelium and regulates NVU functions.

